# Genome Wide Computational Prediction of miRNAs in Kyasanur Forest Disease Virus and their Targeted Genes in Human

**DOI:** 10.1101/095083

**Authors:** Sandeep Saini, Chander Jyoti, Varinder Kumar

**Affiliations:** Department of Bioinformatics, GGDSD College, Sector-32-C, Chandigarh, India, 160031

**Keywords:** Kyasanur forest disease virus, Flavivirus, miRNA, target prediction

## Abstract

RNAs are versatile biomolecules and can be coding or non-coding. Among the non-coding RNAs, miRNAs are small endogenous molecules that play important role in posttranscriptional gene regulation. miRNAs are identified in viruses too and involved in down regulation of host genes. Flavivirus family members are classified in to two groups: mosquito-borne flaviviruses (MBFV) and tick-borne flaviviruses (TBFV). Kyasanur forest disease virus (KFDV) found in India in 1957 (Karnataka) relates to TBFV. Virus has been diffuse to new areas in India and needs attention as it can cause severe hemorrhagic fever. Here in this study, we scanned the virus genome for prediction of miRNAs that can inhibit host target genes. VMir, tool was used for extraction of pre-miRNAs. A total of four miRNAs were found and submitted to ViralMir for classification in to real or pseudo. Interestingly, all four pre-miRNAs were classified as real. Eight mature miRNAs were located in pre-miRNAs by Mature Bayes. A total of 539 human target genes has been identified by using miRDB but ANGPT1 (angiopoietin 1) and TFRC (transferrin receptor) genes were screened to play role in hemorrhagic fever and neurological problems. GO analysis of target genes also supported the evidences.

## Introduction

Kyasanur forest disease virus (KFDV) belongs to family *Flaviviridae*, genus *Flavivirus* [1]. KFDV was first isolated from black faced langur (Presbytis entellus) in Kyasanur forest (Karnataka, India) in March, 1957 [2]. KFDV is mainly transmitted by bite of infected ticks (Haemaphysalis spinigera) and monkeys, rodents and birds acts as reservoir of the virus; hence it is of Zoonotic origin [3]. Upon infection to a human the onset of various symptoms likes fever, frontal headache, myalgia, photophobia, severe prostration, hypotension and hepatomegaly can be commonly seen. But in some severe cases pulmonary, gastrointestinal hemorrhage and neurological manifestations occurred. The term hemorrhagic fever is commonly used for all indications and the disease is termed Kyasanur forest disease (KFD) [4, 5].

Since the first outbreak of KFD in 1957, the disease has been outbreaks many a times from 2012-2015 in its origin state Karnataka and even in new states like Kerala, Tamilnadu and Goa [6, 7]. Serological studies has been confirmed the existence of KFDV in other parts of India including parts of Kutch district, the Saurashtra region in Gujarat state, and in parts of West Bengal state, in forested regions west of Kolkata [8]. An outbreak reported in January, 2016 in Maharashtra has also confirmed 24 KFD cases in human [9]. Number of efforts has been put by the different researchers to make a successful vaccine against KFDV but till date no effective vaccine is available [10–13]. It is evident from fact that even after the administration of vaccine in a population of particular area, the cases has not been stopped [14].

The geographical location of KFDV is not just limited to India, genome analysis studies has shown that it has more than 90 % sequence identity with Alkhurma hemorrhagic fever virus (AHFV), which is isolated in 1994 from blood of hemorrhagic fever patients in Makkah, Saudi Arabia and thought to be transferred by migratory birds from India [15, 16]. Many members of Flaviviridae family such as Dengue [17], Japanese Encephalitis [18], and recently Zika virus [19] were predicted to have encoded miRNAs. It is also evident in past that viral miRNAs can down regulate host genes expression [20–22].

miRNAs are small non-coding RNAs, about 20-25 nt long [23]. In the world of non-coding RNAs, miRNAs are gaining more significance as these small RNA molecules involves in post-transcriptional gene regulation by binding to complementary sites in mRNAs [24]. Experimental determination of miRNA is limited to its expression in specific cell type and time, so the computational approaches are frequently used for prediction of miRNAs [25].

Increase in prevalence of KFDV in India, evolutionary similarity with other viruses found outside India, no effective vaccine against it and evidences of viral miRNAs regulating host gene prompt us to analyze the KFDV genome sequence for miRNA prediction.

## Materials and Methods

### KFDV Genome Sequence

Complete genome sequence of KFDV strain P9605 (Genbank accession number: JF416958.1), a human isolate was obtained from NCBI (National Center for Biotechnology Information). Genome is a single stranded RNA molecule with linear topology and 10774 nucleotide base pairs.

### Pre-miRNAs prediction

Viral genome was scanned for finding self complementary hair pin loop structures of precursor RNAs (pre-miRNAs) by VMir, an ab initio miRNAs prediction program [26]. VMir package contains two individual programs: VMir Analyzer, used for analyzing sequence and VMir Viewer, used for viewing and filtering result of analyzer. The default parameter were used for analyzer (window count: 500, conformation: linear, orientation: both). Stringent filtering was done by setting min. hairpin size: 70, min. score: 115 and min. window count: 35 in VMir viewer as previously described [26].

### Identification of real pre-miRNAs

We used ViralMir, a web interface tool for identification of potential or real viral pre-miRNAs sequences. ViralMir is specially designed for viruses and is based on SVM (support vector machine). The SVM model has been trained on sequence and structural features of experimentally validated pre-miRNAs data set [27].

### Secondary structure prediction and energy calculation

The Mfold web server with default parameters was used to predict the secondary structure and minimum free energy (MFE) of pre miRNAs [28].

### Extraction of mature miRNAs from pre-miRNAs

For further downstream analysis, mature miRNAs were extracted from pre-miRNAs sequences using Mature Bayes- an online tool that uses Naive Bayes Classifier (NBC) taking into account sequence as well as structural information of experimental predicted miRNA precursors [29].

### Target gene prediction by miRDB

miRDB, a web based server which provide custom target prediction to user to identify the target gene in human was used. All the mature miRNAs was submitted individually for identification of target genes. miRDB uses the seeding approach and scan the 3’ UTR (untranslated regions) of human’s gene for possible hybridization with miRNAs sequence [30].

### Literature data mining and Screening of genes

NCBI Gene database was used for retrieving the list of genes (supplementary data Table 1) involved in hemorrhagic fever. Manual crosschecking was done with target genes retrieved from miRDB. Literature search was carried out for supporting the screened gene result.

### GO (Gene Ontology) analysis

Most of the miRNAs prediction works are limited to up to target prediction but now-a-days gene ontology studies were being done to gain insight in to molecular functional, biological process and cellular component of the target genes [31]. We also used PANTHER for analyzing our entire predicted target gene for GO terms [32, 33]. NCBI gene IDs of target genes in a text files were used for this analysis.

## Results and Discussion

### Prediction of pre-miRNAs in KFDV

An Ab-intio based program analyzes the sequence feature for making any prediction [34–36]. Genome organization studies of miRNAs have revealed that they can be present on introns of protein-coding transcriptional unit (TU), introns of non-coding TU, exons of non-coding TU and exons of coding TU [37–40]. On whatever location they may present but they tend to stabilize themselves by complementary hybridization to form hairpin loop structural features [41, 42]. Our study is also based on above mentioned concept.

After analysis of complete genome of KFDV accession number JF416958.1 by VMir we got four pre-miRNAs (Fig. 1) at stringent parameters described above in methodology. Out of four miRNAs three were found in reverse strand and one on direct strand. All the predicted miRNAs were in size range 147-185 nt. The genomic position of each pre-miRNAs varies and is shown in Table 1 along with VMir score.

**Figure 1:**
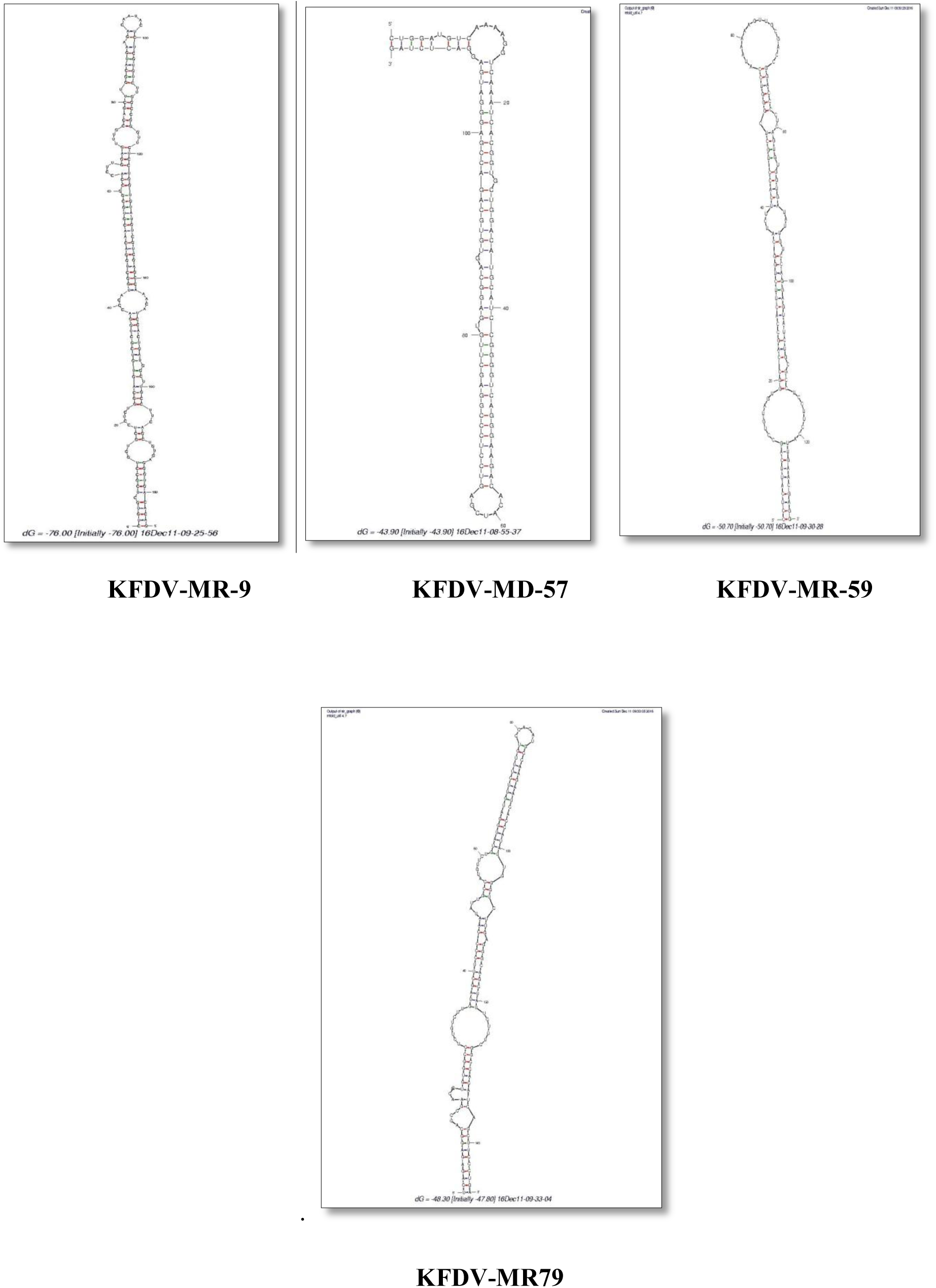
Pre-miRNAs resulted by Mfold

**Table 1.** Predicted pre-miRNAs in KFDV genome

| Sr No. | Predicted pre-miRNA | RANK | STRAND ORIENTATION | Size pre-miRNA(nt) | Position on Genome | VMIR Score |
| --- | --- | --- | --- | --- | --- | --- |
| 1 | KFDV-MR9 | 1 | Reverse | 185 | 1061 - 1245 | 200.6 |
| 2 | KFDV-MD57 | 2 | Direct | 116 | 5165 - 5280 | 193.3 |
| 3 | KFDV-MR59 | 3 | Reverse | 131 | 6062 - 6192 | 172.3 |
| 4 | KFDV-MR79 | 4 | Reverse | 147 | 8488 - 8634 | 120.2 |

All the four pre-miRNAs were identified as real by virus specific tool, ViralMir and their minimum free energy calculated by Mfold is shown in Table 2.

**Table 2.** Classification of pre-miRNAs and MFEs

| Sr No. | Predicted pre-miRNA | REAL\PSEUDO | pre-miRNA MFEs (-$\Delta G$. kCAL/MoL) |
| --- | --- | --- | --- |
| 1 | KFDV-MR9 | REAL | $\Delta G = -76.00$ kcal/mol |
| 2 | KFDV-MD57 | REAL | $\Delta G = -43.90$ kcal/mol |
| 3 | KFDV-MR59 | REAL | $\Delta G = -50.70$ kcal/mol |
| 4 | KFDV-MR79 | REAL | $\Delta G = -47.80$ kcal/mol |

### Mature miRNAs extraction

Pre-miRNAs are not functional mature miRNAs, Dicer cleaved pre-miRNAs in to small ∼22 nt long mature miRNAs molecules. After processing large pre-miRNAs, two single strands one in 5’ stem and other in 3’ stem were obtained. One or both strands can serve as mature miRNA molecule depending on the assembly of RISC complex [43, 44]. We predicted mature miRNAs from each four pre-miRNAs both in 5’ stem and 3’ stem by Mature Bayes tool. A total of eight mature miRNAs were predicted and their sequence is shown in Table 3.

**Table 3.**
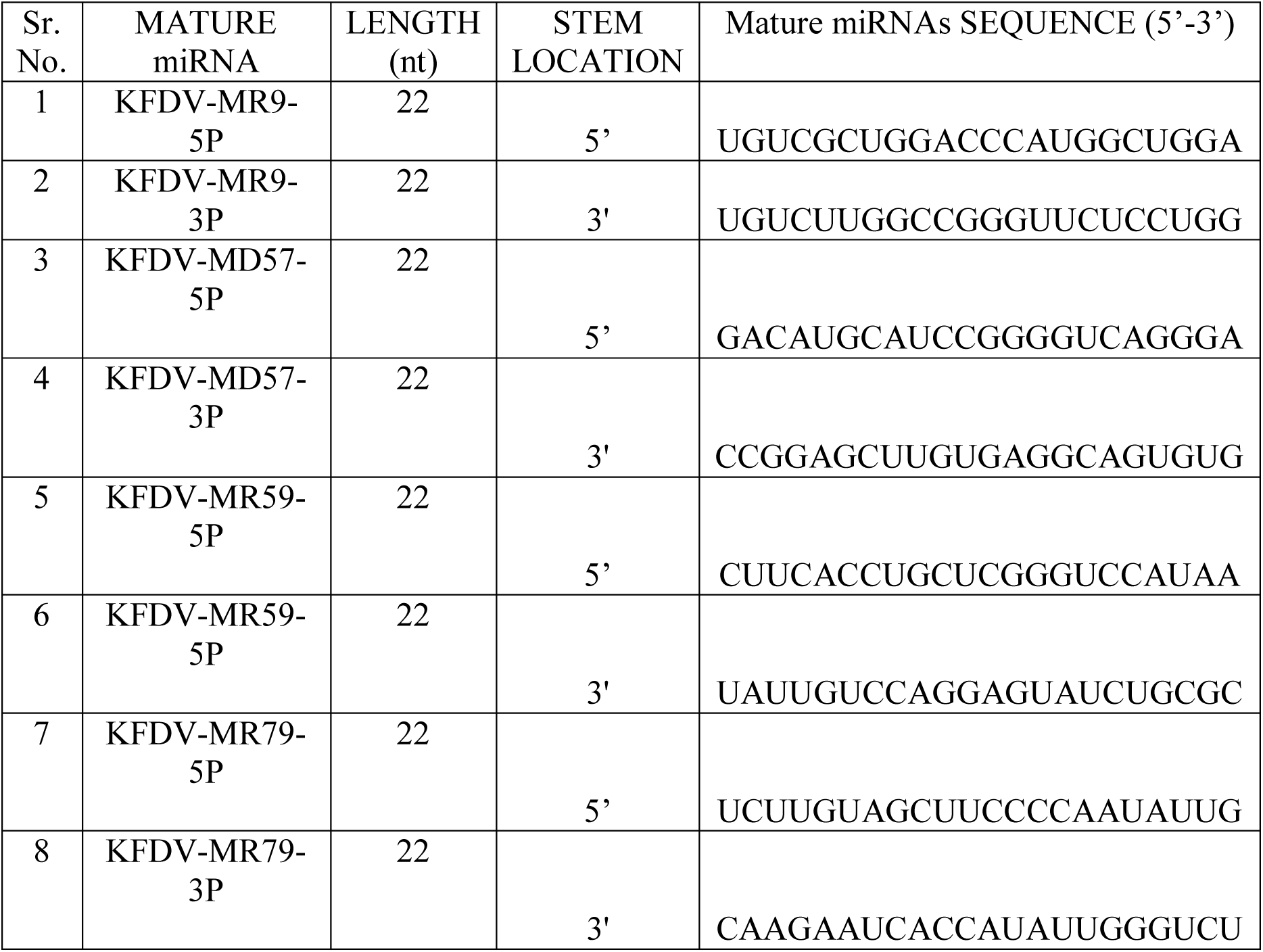
Mature miRNAs

### Prediction of target genes in human

miRNAs target prediction can be done by experimentally or computationally. Where the experimental evidence is hard to obtained, computational predictions mainly relies on the Watson-Crick base pairing between miRNA and mRNA molecule. Most of the algorithm used seed pairing approach for predicting interaction between miRNA and “seed sites” on mRNA molecule [45–47].

To follow this approach, we also used miRDB server for prediction of target gene in human. The server uses the MirTarget algorithm, which is based on 7-mer seeding approach and custom predict miRNAs targets in human gene’s 3’ UTRs [48]. We successfully predicted 539 target genes with miRDB score >80, shown in supplementary data table 2. Crosschecking with NCBI gene list for hemorrhagic fever gave only two genes, ANGPT1 (angiopoietin 1) and TFRC (transferrin receptor). Both ANGPT1 and TFRC play role in blood vessel stability and neurological development respectively. PubMed ids of literature evidences of these genes were given in table 4.

**Table 4.** PubMed ids of literature evidence

| Mature miRNA | Target Gene | Description | Reference PMID |
| --- | --- | --- | --- |
| KFDV-MD57-5P | ANGPT1 | angiopoietin 1 | 20156556,<br>16645151 |
|  | TFRC | transferrin<br>receptor | 26339443,<br>19211831 |

### GO analysis

Target genes were found to be involved in a number of pathways (Fig. 2) including synaptic vesicles trafficking pathway (P05734), axon guidance (P00008) that play important role in proper nervous system functioning, pathways like coagulation (P00011), plasminogen activating cascade (P00050), angiogenesis (P00005) confirmed role in blood regulation. Whereas involvement of T cell activation pathway (P00053) can be related to breach in the immune system. Biological processes (Fig. 3) involvements were also spread to immune system (GO:0002376), response to stimulus (GO:0050896) and biological adhesion (GO:0022610) etc. Molecular functions (Fig. 4) of the targeted genes is found to be involved in transporter activity (GO:0005215), translation regulator activity (GO:0045182), binding activity (GO:0005488) and signal transducer activity (GO:0004871) etc. Cellular component analysis (Fig. 5) revealed that target gene products were the part of synapses (GO:0045202), extracellular matrix (GO:0031012) and membrane (GO:0016020) etc.

**Figure 2:**
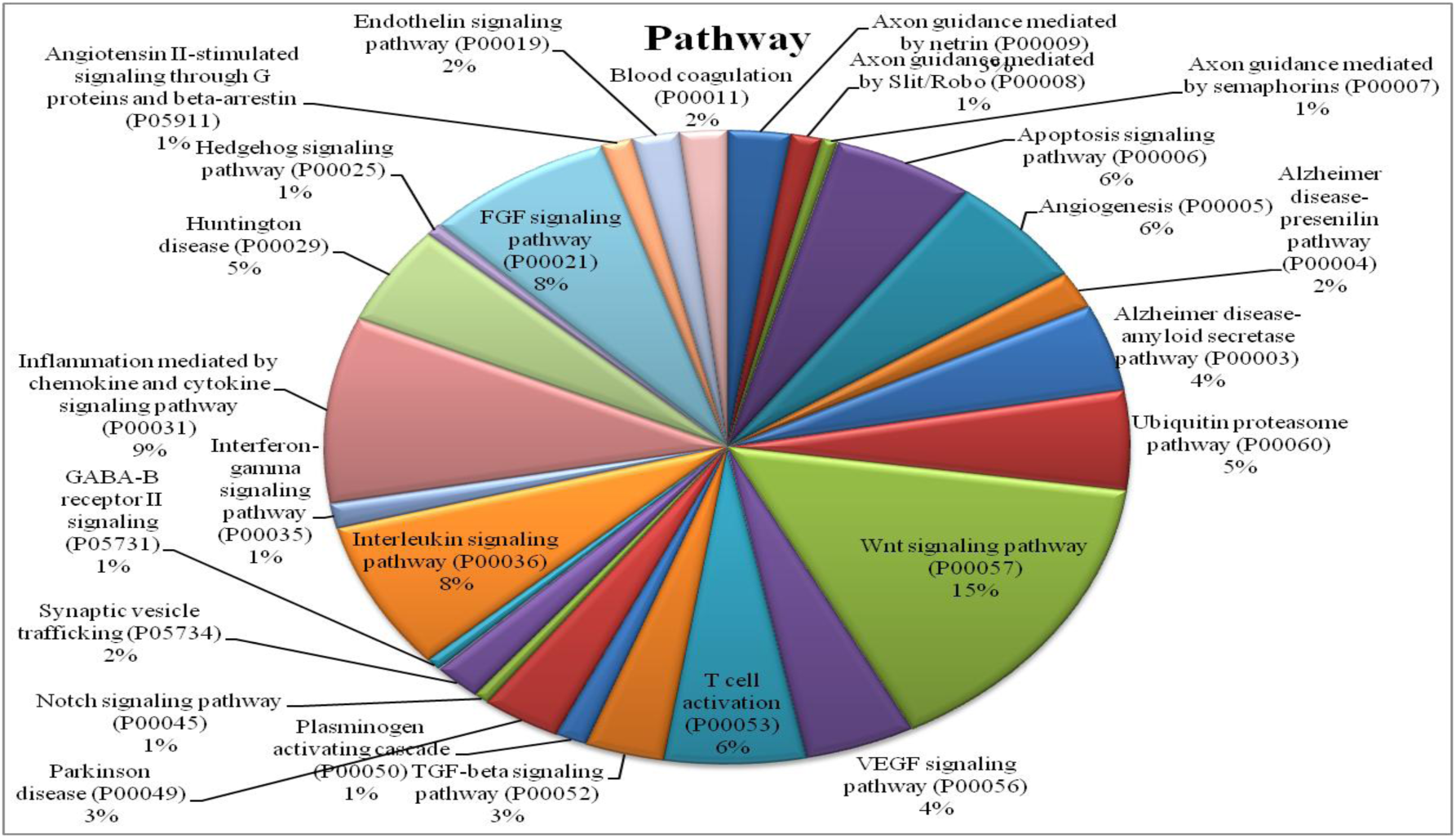
Pathway analysis of target genes

**Figure: 3:**
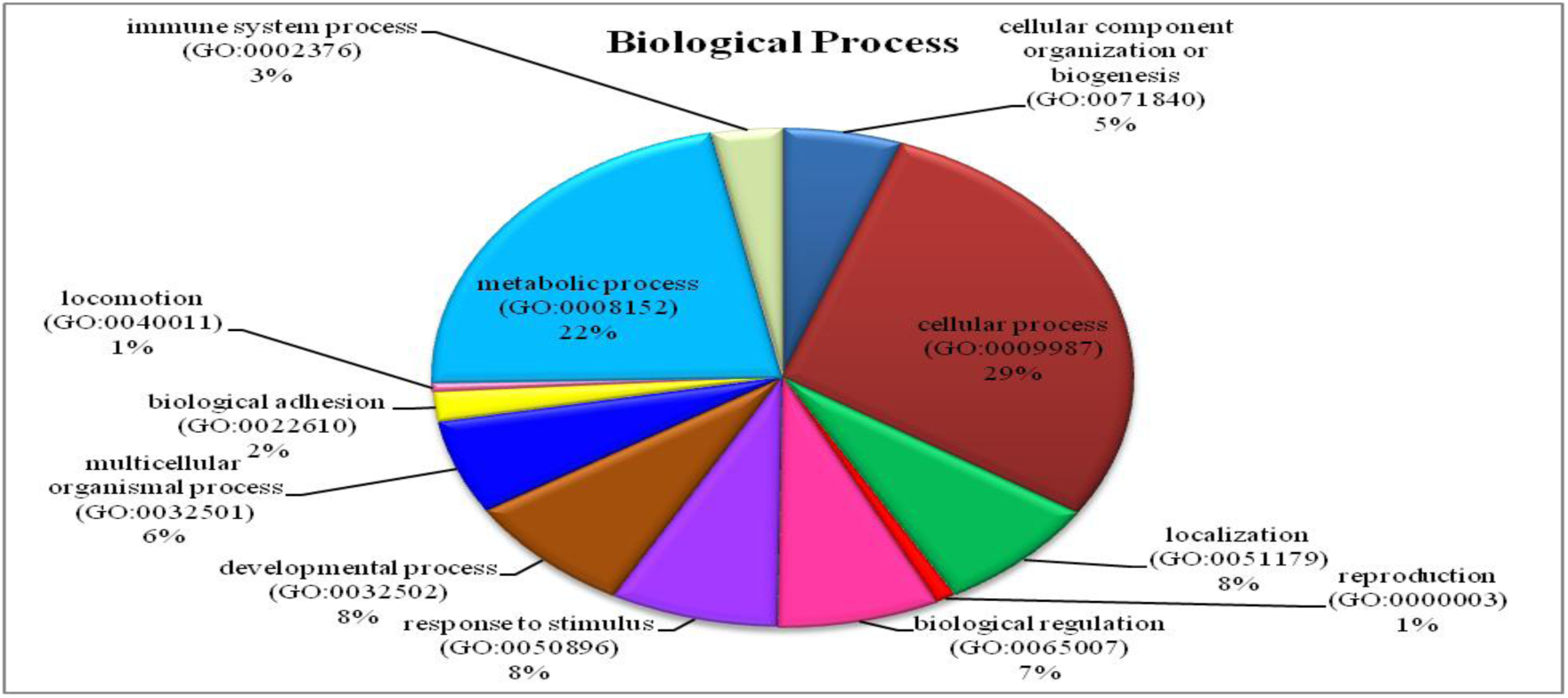
Biological Process of target genes

**Figure 4:**
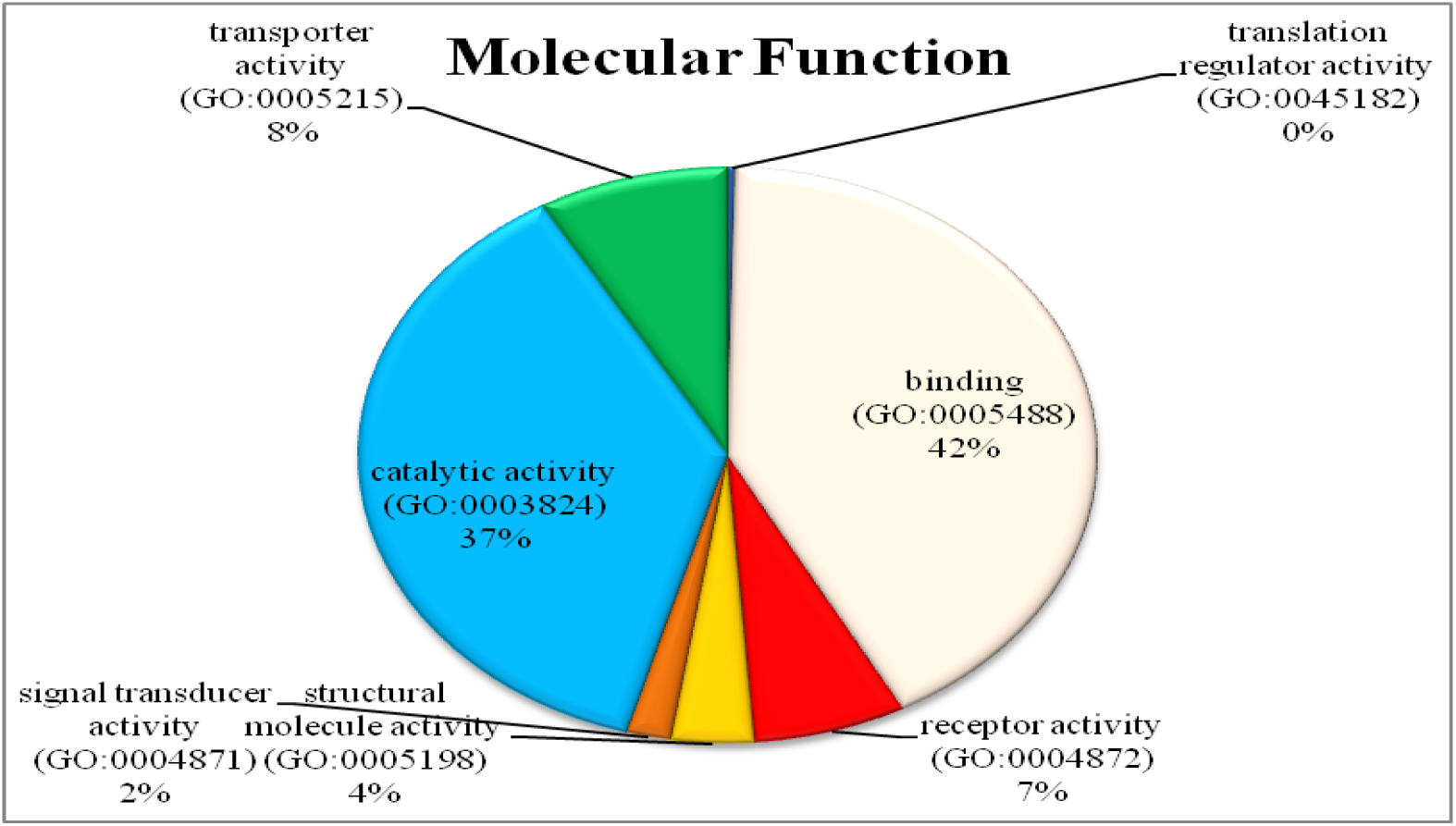
Molecular functional analysis of target genes

**Figure 5:**
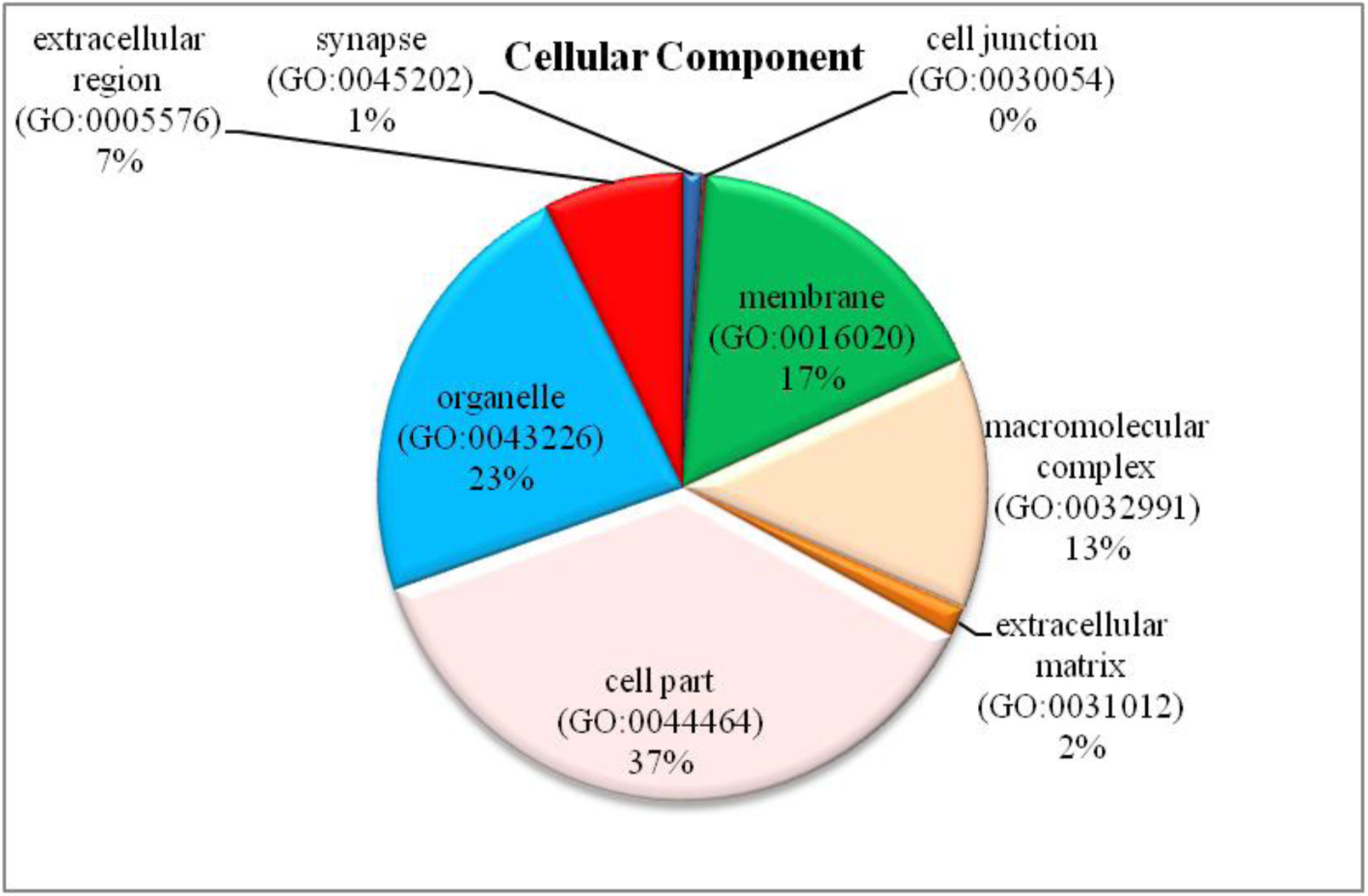
Cellular component analysis of target genes

### Conclusion

Ebola and Zika virus outbreak in 2014 [49] and 2015 [50] respectively posed a serious problem to world. Kyasanur forest disease virus (KFDV) in India has been outbreak time to time, caused hemorrhagic fever and neurological disturbances [51]. In our study, we have predicted eight mature miRNAs in KFDV genome and their target genes in human. Target gene analysis after data mining and literature support had screen out two genes, ANGPT1 (angiopoietin 1) and TFRC (transferrin receptor). ANGPT1 encodes a secreted glycoprotein, which plays a critical role in reciprocal interaction between endothelium and surrounding matrix, inhibits endothelial permeability and contributes to blood vessel maturation and stability. TFRC encodes a cell surface receptor necessary for cellular iron uptake. This receptor is required for erythropoiesis and neurological development. Both these target genes were predicted to have hybridized with KFDV-MD57-5P mature miRNA.

In addition, gene ontology analysis has predicted pathways related to blood coagulation, angiogenesis and axon guidance. Biological processes were related to immune system, response to stimulus and biological adhesion. Cellular processes were involved in synapses. To summarize with, the final two predicted target genes ANGPT1 and TFRC molecular function relates to Kyasanur forest disease etiology. Our work involved in-silico predictions in combination with literature mining to support for predictions results. But it needs to be still verifying by in-vitro designing of these predicted miRNAs constructs and their hybridization with the proposed target genes.

## References

1. Pattnaik P. Kyasanur forest disease: an epidemiological view in India. Rev Med Virol. 2006 May-Jun; 16(3):151–65.

2. Work T. H., Roderiguez F. R., Bhatt P. N. Virological Epidemiology of the 1958 Epidemic of Kyasanur Forest Disease. Am J Public Health Nations Health. 1959 Jul; 49(7): 869–874.

3. Roy P, Maiti D, Goel MK, Rasania S. Kyasanur Forest Disease: An emerging tropical disease in India. J Res Med Den Sci. 2014; 2(2): 1–4. doi:10.5455/jrmds.2014221

4. Adhikari Prabha MR, Prabhu MG, Raghuveer CV, Bai M, Mala MA. Clinical study of 100 cases of Kyasanur Forest disease with clinicopathological correlation. Indian J Med Sci. 1993 May; 47(5): 124–30.

5. Pavri K. Clinical, clinicopathologic, and hematologic features of Kyasanur Forest disease. Rev Infect Dis. 1989 May-Jun; 11 Suppl 4:S854–9.

6. Murhekar MV, Kasabi GS, Mehendale SM, Mourya DT, Yadav PD, Tandale BV. On the transmission pattern of Kyasanur Forest disease (KFD) in India. Infect Dis Poverty. 2015 Aug 19; 4:37. doi: 10.1186/s40249-015-0066-9.

7. Yadav PD, Shete AM, Patil DY, Sandhya VK, Prakash KS, Surgihalli R, Mourya DT. Outbreak of Kyasanur Forest disease in Thirthahalli, Karnataka, India, 2014. Int J Infect Dis. 2014 Sep; 26:132–4. doi: 10.1016/j.ijid.2014.05.013. Epub 2014 Jul 22.

8. Holbrook MR. Kyasanur Forest Disease. Antiviral Res. 2012 Dec; 96(3): 353–362. Published online 2012 Oct 27. doi: 10.1016/j.antiviral.2012.10.005.

9. Awate P, Yadav P, Patil D, Shete A, Kumar V, Kore P, Dolare J, Deshpande M, Bagde S, Sapkal G, Gurav Y, Mourya DT. Outbreak of Kyasanur Forest disease (monkey fever) in Sindhudurg, Maharashtra State, India, 2016. J Infect. 2016 Jun;72(6):759–61. doi: 10.1016/j.jinf.2016.03.006. Epub 2016 Mar 18.

10. Kasabi GS, Murhekar MV, Sandhya VK, Raghunandan R, Kiran SK, Channabasappa GH, Mehendale SM. Coverage and effectiveness of Kyasanur forest disease (KFD) vaccine in Karnataka, South India, 2005-10. PLoS Negl Trop Dis. 2013;7(1):e2025. doi: 10.1371/journal.pntd.0002025. Epub 2013 Jan 24.

11. Kiran SK, Pasi A, Kumar S, Kasabi GS, Gujjarappa P, Shrivastava A, Mehendale S, Chauhan LS, Laserson KF, Murhekar M. Kyasanur Forest disease outbreak and vaccination strategy,Shimoga District, India, 2013-2014. Emerg Infect Dis. 2015 Jan;21(1):146–9. doi: 10.3201/eid2101.141227.

12. Shah KV, Dandawate CN, Bhatt PN. Kyasanur forest disease virus: viremia and challenge studies in monkeys with evidence of cross-protection by Langat virus infection. F1000Res. 2012 Dec 7;1:61. doi: 10.12688/f1000research.1-61.v1. eCollection 2012.

13. Cook BW, Ranadheera C, Nikiforuk AM, Cutts TA, Kobasa D, Court DA, Theriault SS. Limited Effects of Type I Interferons on Kyasanur Forest Disease Virus in Cell Culture. PLoS Negl Trop Dis. 2016 Aug 1;10(8):e0004871. doi: 10.1371/journal.pntd.0004871. eCollection 2016.

14. Dandawate CN, Upadhyaya S, Banerjee K. Serological response to formolized Kyasanur forest disease virus vaccine in humans at Sagar and Sorab Talukas of Shimoga district. J Biol Stand. 1980;8(1):1–6.

15. Dodd KA, Bird BH, Khristova ML, Albariño CG, Carroll SA, Comer JA, Erickson BR, Rollin PE, Nichol ST. Ancient ancestry of KFDV and AHFV revealed by complete genome analyses of viruses isolated from ticks and mammalian hosts. PLoS Negl Trop Dis. 2011 Oct;5(10):e1352. doi: 10.1371/journal.pntd.0001352. Epub 2011 Oct 4.

16. Charrel RN, Zaki AM, Attoui H, Fakeeh M, Billoir F, Yousef AI, de Chesse R, De Micco P, Gould EA, de Lamballerie X. Complete coding sequence of the Alkhurma virus, a tick-borne flavivirus causing severe hemorrhagic fever in humans in Saudi Arabia. Biochem Biophys Res Commun. 2001 Sep 21;287(2):455–61.

17. Ospina-Bedoya M, Campillo-Pedroza N, Franco-Salazar JP, Gallego-Gomez JC. Computational Identification of Dengue Virus MicroRNA-Like Structures and their Cellular Targets. Bioinform Biol Insights. 2014 Aug 14;8:169–76. doi: 10.4137/BBI.S13649. eCollection 2014.

18. Vijay Laxmi S, Alka D. In silico identification of miRNAs and their target prediction from Japanese encephalitis J. Bioinform. Seq. Anal. 2013 Feb; 5(2); 25–33.

19. Pylro VS, Oliveira FS, Morais DK, Cuadros-Orellana S, Pais FS, Medeiros JD, Geraldo JA, Gilbert J, Volpini AC, Fernandes GR. ZIKV - CDB: A Collaborative Database to Guide Research Linking SncRNAs and ZIKA Virus Disease Symptoms. PLoS Negl Trop Dis. 2016 Jun 22;10(6):e0004817. doi: 10.1371/journal.pntd.0004817. eCollection 2016.

20. Carl JW, Trgovcich J, Hannenhalli S. Widespread evidence of viral miRNAs targeting host pathways. BMC Bioinformatics. 2013;14:S3.

21. Grundhoff A, Sullivan CS. Virus-encoded microRNAs. Virology. 2011 Mar 15;411(2):325–43. doi: 10.1016/j.virol.2011.01.002. Epub 2011 Jan 31.

22. Skalsky RL, Cullen BR. Viruses, microRNAs, and host interactions. Annu Rev Microbiol. 2010; 64:123–41. doi: 10.1146/annurev.micro.112408.134243.

23. Grosshans H, Filipowicz W. Molecular biology: The expanding world of small RNAs. Nature. 2008 Jan 24;451(7177):414–6. doi: 10.1038/451414a.

24. He L, Hannon GJ. MicroRNAs: small RNAs with a big role in gene regulation. Nat Rev Genet. 2004 Jul; 5(7):522–31.

25. Bartel DP. MicroRNAs: genomics, biogenesis, mechanism, and function. Cell. 2004 Jan 23; 116(2):281–97.

26. Grundhoff A1, Sullivan CS, Ganem D. A combined computational and microarray-based approach identifies novel microRNAs encoded by human gamma-herpesviruses. RNA. 2006 May; 12(5):733–50. Epub 2006 Mar 15.

27. Huang KY, Lee TY, Teng YC, Chang TH. ViralmiR: a support-vector-machine-based method for predicting viral microRNA precursors. BMC Bioinformatics. 2015;16 Suppl 1:S9. doi: 10.1186/1471-2105-16-S1-S9. Epub 2015 Jan 21.

28. Michael Z. Mfold web server for nucleic acid folding and hybridization prediction. Nucleic Acids Res. 2003 Jul 1; 31(13): 3406–3415.

29. Gkirtzou K., Tsamardinos I., Tsakalides P. and Poirazi P. MatureBayes: a probabilistic algorithm for identifying the mature miRNA within novel precursors. PLoS One 5, e11843, doi: (2010).10.1371/journal.pone.0011843.

30. Wong N, Wang X. miRDB: an online resource for microRNA target prediction and functional annotations. Nucleic Acids Res. 2015 Jan;43(Database issue):D146–52. doi: 10.1093/nar/gku1104. Epub 2014 Nov 5.

31. Teng Y, Wang Y, Zhang X, Liu W, Fan H, Yao H, Lin B, Zhu P, Yuan W, Tong Y, Cao W. Systematic Genome-wide Screening and Prediction of microRNAs in EBOV During the 2014 Ebolavirus Outbreak. Sci Rep. 2015 May 26;5:9912. doi: 10.1038/srep09912

32. Gene Ontology Consortium: going forward. Nucleic Acids Res. 2015 Jan;43(Database issue):D1049–56. doi: 10.1093/nar/gku1179. Epub 2014 Nov 26.

33. Mi H, Muruganujan A, Casagrande JT, Thomas PD. Large-scale gene function analysis with the PANTHER classification system. Nat Protoc. 2013 Aug;8(8):1551–66. doi: 10.1038/nprot.2013.092. Epub 2013 Jul 18.

34. Allmer J, Yousef M. Computational methods for ab initio detection of microRNAs. Front Genet. 2012 Oct 10;3:209. doi: 10.3389/fgene.2012.00209. eCollection 2012.

35. Kumar S, Ansari FA, Scaria V. Prediction of viral microRNA precursors based on human microRNA precursor sequence and structural features. Virol J. 2009 Aug 20;6:129. doi: 10.1186/1743-422X-6-129.

36. Tempel S, Tahi F. A fast ab-initio method for predicting miRNA precursors in genomes. Nucleic Acids Res. 2012 Jun;40(11):e80. doi: 10.1093/nar/gks146. Epub 2012 Feb 22.

37. Kim VN, Han J, Siomi MC. Biogenesis of small RNAs in animals. Nat Rev Mol Cell Biol. 2009 Feb;10(2):126–39. doi: 10.1038/nrm2632.

38. Kim VN, Nam JW. Genomics of microRNA. Trends Genet. 2006 Mar;22(3):165–73. Epub 2006 Jan 30.

39. Bavia L, Mosimann AL, Aoki MN, Duarte Dos Santos CN. A glance at subgenomic flavivirus RNAs and microRNAs in flavivirus infections. Virol J. 2016 May 28;13:84. doi: 10.1186/s12985-016-0541-3.

40. Olena AF, Patton JG. Genomic organization of microRNAs. J Cell Physiol. 2010 Mar;222(3):540–5. doi: 10.1002/jcp.21993.

41. Berezikov E. Evolution of microRNA diversity and regulation in animals. Nat Rev Genet. 2011 Nov 18;12(12):846–60. doi: 10.1038/nrg3079.

42. Varble A, Chua MA, Perez JT, Manicassamy B, García-Sastre A, tenOever BR. Engineered RNA viral synthesis of microRNAs. Proc Natl Acad Sci U S A. 2010 Jun 22;107(25):11519–24. doi: 10.1073/pnas.1003115107. Epub 2010 Jun 7.

43. Nelson P, Kiriakidou M, Sharma A, Maniataki E, Mourelatos Z. The microRNA world: small is mighty. Trends Biochem Sci. 2003 Oct;28(10):534–40.

44. Li Guo and Zuhong Lu. The Fate of miRNA* Strand through Evolutionary Analysis: Implication for Degradation As Merely Carrier Strand or Potential Regulatory Molecule? PLoS One. 2010; 5(6): e11387. Published online 2010 Jun 30. doi: 10.1371/journal.pone.0011387.

45. Rajewsky N. microRNA target predictions in animals. Nat Genet. 2006 Jun;38 Suppl:S8–13.

46. Saito T, Saetrom P. MicroRNAs--targeting and target prediction. N Biotechnol. 2010 Jul 31;27(3):243–9. doi: 10.1016/j.nbt.2010.02.016. Epub 2010 Feb 26.

47. Akbari Moqadam F1, Pieters R, den Boer ML. The hunting of targets: challenge in miRNA research. Leukemia. 2013 Jan;27(1):16–23. doi: 10.1038/leu.2012.179. Epub 2012 Jul 3.

48. Wang X. Improving microRNA target prediction by modeling with unambiguously identified microRNA-target pairs from CLIP-ligation studies. Bioinformatics. 2016 May 1;32(9):1316–22. doi: 10.1093/bioinformatics/btw002. Epub 2016 Jan 6.

49. Cuevas EL, Tong VT, Rozo N, Valencia D, Pacheco O, Gilboa SM, Mercado M, Renquist CM, González M, Ailes EC, Duarte C, Godoshian V, Sancken CL, Turca AM, Calles DL, Ayala M, Morgan P, Perez EN, Bonilla HQ, Gomez RC, Estupiñan AC, Gunturiz ML, Meaney-Delman D, Jamieson DJ, Honein MA, Martínez ML. Preliminary Report of Microcephaly Potentially Associated with Zika Virus Infection During Pregnancy - Colombia, January-November 2016. MMWR Morb Mortal Wkly Rep. 2016 Dec 16;65(49):1409–1413. doi: 10.15585/mmwr.mm6549e1.

50. Pavot V. Ebola virus vaccines: Where do we stand? Clin Immunol. 2016 Dec;173:44–49. doi: 10.1016/j.clim.2016.10.016. Epub 2016 Oct 28.

51. John JK, Kattoor JJ, Nair AR, Bharathan AP, Valsala R, Sadanandan GV. Kyasanur forest disease: a status update. Adv. Anim. Vet. Sci. 2014 July; 2 (6): 329–336.

